# Germline VCF Annotator: a lightweight pipeline for processing germline VCFs with robust variant extraction and read evidence quality control

**DOI:** 10.64898/2026.04.06.716730

**Authors:** Zarko Manojlovic

**Author notes:** Correspondence should be addressed to Z.M.

## Abstract

Raw variant calls are typically distributed as VCF files and are not well-suited for direct human review. They are intended for programmatic parsing, and spreadsheet import can distort data through automatic type conversion. Furthermore, variants in VCF are commonly annotated to add gene context and predicted functional consequences. Ensembl VEP, a widely used standard for transcript-aware variant annotation, was adapted in this study to generate standardized consequence fields across genomic features. Using a colon crypt whole-genome sequencing cohort as the motivating dataset, this study examined whether variation at DNA damage response and repair (DDR) loci could contribute to mutation-burden patterns in normal colon crypts, including patterns associated with age and potential treatment-related exposure. To make this question testable in a reproducible table-based format, the Germline VCF Annotator was developed as a two-step workflow that normalizes germline VCFs, generates VEP tabular annotations with explicit allele fields, and then extracts variants of interest and appends read-evidence metrics to assign a rules-based QC class. Within-patient concordance across technical repeats at predefined DDR loci was near-perfect after filtering for nonsilent SNVs with read depth ≥15, with discordance concentrated among Low-QC loci. Bulk and crypt-derived samples showed no age-related trend in DDR burden. Although the demonstration centers on DDR and aging, the Germline VCF Annotator is applicable to other gene sets that require human-readable locus-level summaries with retained allele provenance and read evidence.

## INTRODUCTION

High-throughput sequencing has made germline variant calling routine, but transforming a raw VCF into an interpretable, analysis-ready table remains far from standardized. The Variant Call Format (VCF) is an excellent interchange standard, yet it is optimized for compact representation and machine parsing. As a result, basic interpretive questions, such as which variant is present, what consequences it has, and how well supported the call is, are often not immediately accessible. This burden is commonly shifted downstream to custom scripts and spreadsheet-based review, which increases the risk of inconsistency, particularly for allele representation, multiallelic sites, and transcript-dependent consequences^1–3^.

Annotation is a critical step in making variants biologically interpretable and can include consequence assignment, genomic context mapping, and integration of external resources such as population frequency and clinical databases. For coding and splice relevant small variants, several established effect annotators, including Ensembl VEP and SnpEff, are transcript-aware and can assign distinct consequences to the same genomic variant across overlapping transcripts and genomic features^4,5^. Ensembl VEP, a widely used annotator, provides standardized consequence assignments, supports commonly used transcript models including Ensembl and RefSeq, and offers flexible tabular output well suited for downstream extraction and restructuring^4,6–8^. Furthermore, VEP was adapted to generate annotations in a format suitable for allele provenance tracking, concordance assessment, and reproducible locus-level reporting. However, because transcript-aware annotation can assign multiple consequences to a single genomic event, a single locus may expand into multiple annotation rows. Although this is biologically appropriate, it complicates unique variant counting, concordance assessment, and consistent reporting^5,9–11^.

In somatic genomics, a tool such as vcf2maf became widely adopted because it converts annotated variants into standardized human-readable tables and applies consistent transcript handling for downstream filtering, visualization, and reporting^12^. Germline workflows more often stop at VCF plus annotation, leaving investigators to assemble the reporting layer themselves. This limitation becomes especially apparent in studies that require transparent quality review, replicate concordance, and collaborator-facing summaries without requiring every user to work directly from VCF internals. Unrelated utilities with similar names such as vcf2csv (https://vcf2csv.sourceforge.net/) exist for converting vCard .vcf contact files to CSV or HTML, but these do not operate on genomic VCF files and are not designed for variant annotation and review.

Normal human colon crypts provide a well-defined clonal unit for studying somatic mutation accumulation in aging tissue. Prior work in normal colorectal epithelium has shown that somatic mutation burden increases with age and that individual crypts can display substantial mutational heterogeneity, making this tissue a useful system for resolving background mutation processes at single crypt resolution^13–16^. Within that setting, accurate interpretation of somatic patterns also requires careful definition of the inherited germline background. In particular, constitutional variations in genes involved in the DNA damage response and repair can influence mutation accumulation, complicating crypt-level comparisons or creating the appearance of biologically meaningful differences that are instead explained by inherited background. The immediate motivation for developing the Germline VCF Annotator arose from whole-genome sequencing studies of normal human colon crypts^15–17^. The practical need in these colon crypt studies was not only to annotate germline variants, but also to generate gene focused tables that remained true to the underlying evidence and reproducible across repeated cohort scale analyses.

To provide a biologically interpretable demonstration and enable consistent cross sample comparison, downstream analyses focused on a predefined set of DNA damage response and repair (DDR) genes. DDR genes function in pathways that detect, signal, and resolve DNA lesions, including mismatch repair, base excision repair, nucleotide excision repair, homologous recombination, nonhomologous end joining, and Fanconi anemia associated repair^18,19^. This focus was motivated by the question of whether germline variations in DDR pathways lead to age associated differences in mutation burden in normal colon crypts, and whether occult DDR variants in otherwise healthy individuals could inflate mutation burden patterns. The DDR gene set, therefore, provides a practical framework for evaluating concordance, quality behavior, and annotation-driven reporting in germline variant analysis while anchoring the workflow to a biologically meaningful hypothesis.

## MATERIALS AND METHODS

### Access

Germline VCF Annotator is publicly available (source code) on GitHub (https://github.com/Z-Mano/Germline-VCF-Annotator), including the complete two-step workflow scripts, documentation, and example configuration files. The software is licensed for academic and research use only (© 2026 Zarko Manojlovic and Chih-Lin Hsieh).

### Workflow overview

Germline VCF Annotator is organized as a two-step workflow (Fig. 1) that converts germline VCF outputs into analysis-ready tables while preserving allele identity and read evidence provenance. For this study, a curated DNA damage response and repair (DDR) gene set (Supplemental Table S1) defined the targeted interval space for downstream extraction, QC stratification, concordance analysis, and ClinVar-focused review. Although the workflow was demonstrated here using FreeBayes germline VCFs, the downstream normalization, annotation, and extraction framework is broadly compatible with germline VCFs produced by other commonly used callers (e.g., GATK HaplotypeCaller and DeepVariant), provided that the standard VCF structure and the required evidence fields are retained.

**Fig. 1.**
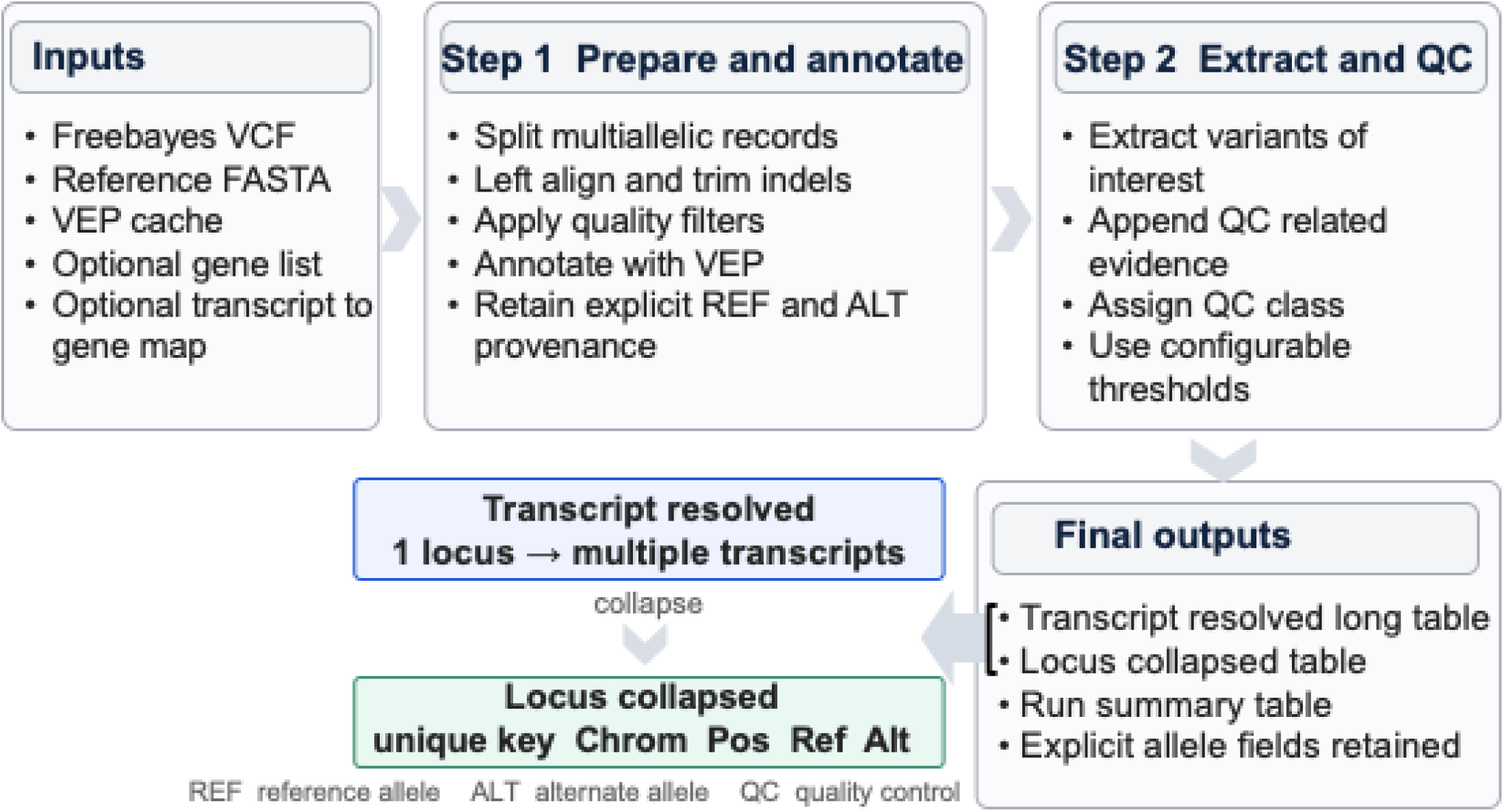
Germline VCF Annotator workflow overview. The figure summarizes the two-step workflow and the relationship between transcript resolved and locus collapsed outputs. In Step 1, input germline VCFs are prepared and normalized, followed by Ensembl VEP annotation to generate transcript resolved tabular output with explicit reference and alternate allele provenance. In Step 2, variants of interest are extracted, QC related metrics from the VCF are appended, and an adjustable QC class is assigned. Final outputs include a transcript resolved long table, a locus collapsed table keyed by chromosome, position, reference allele, and alternate allele, and a run-level summary. The transcript resolved view preserves transcript specific variant annotations and mechanistic context, whereas the locus collapsed view provides a stable genomic representation for burden counting, concordance analysis, and site level review.

### Required inputs

At minimum, the pipeline requires a germline VCF, either single sample or multi sample, preferably bgzipped and indexed, a local Ensembl VEP installation with offline cache recommended, and a reference genome FASTA consistent with the VCF build, for example, GRCh38. Optional inputs include a gene list for targeted reporting and a transcript to gene mapping table to harmonize transcript identifiers across resources.

### Step 1. VCF normalization and VEP annotation

The first stage prepares germline VCFs for stable downstream parsing by applying standard VCF normalization followed by Ensembl VEP annotation. In this study, normalization refers to canonicalizing variant representation so that the same event is encoded consistently across variant outputs. This includes decomposing multiallelic records into one reference and one alternate event per record, along with left alignment and trimming of indels against the reference genome. These steps generate stable locus identifiers defined by chromosome, position, reference allele, and alternate allele, which are then used to collapse and compare concordance. Following normalization, Ensembl VEP together with bcftools (SAMtools suites)^1,20^ was used to generate a tab-delimited annotation file that preserves transcript-resolved consequences while explicitly retaining allele provenance, including the reference allele and the uploaded allele, thereby avoiding downstream ambiguity. Output artifacts from this stage typically include normalized VCFs and VEP-annotated TSV files containing consequence annotations and explicit allele fields.

### Step 2. Variant extraction and read evidence QC

The second stage parses the VEP TSV and combines it with read-level evidence from the input VCF to generate human-readable CSV tables (Fig. 1). Variants can be extracted globally or restricted to a user-defined gene set. Transcript resolved rows are preserved so that each transcript consequence remains explicit. In parallel, a locus collapsed representation is generated by aggregating records by genomic coordinates and alleles. Read evidence metrics are then appended to assign an evidence-based QC class derived from depth, mapping quality, strand balance, and related signals. This enables reproducible filtering and more transparent interpretation. Output artifacts include a transcript resolved long table, a locus collapsed table, and a run level summary table.

### Output interpretation

The long table is most useful when the transcript context is central to interpretation, including transcript selection, HGVSc and HGVSp review, and comparison of transcript-specific consequences. The collapsed table is the preferred representation for locus level burden, concordance metrics across technical repeats, and downstream analyses that treat each genomic site as a single event.

Output files include:

- germline_<…>_with_qc_<run_id>.csv (transcript-resolved long table)
- germline_<…>_with_qc_collapsed_<run_id>.csv (locus-collapsed table)
- germline_<…>_with_qc_summary_<run_id>.csv (run-level summary and counts)

### Quality control and evidence grading

A central goal of Germline VCF Annotator is to make technical confidence directly interpretable in spreadsheet-compatible output without discarding the underlying read evidence. To achieve this, the pipeline derives a rule-based QC class for each variant from read-level metrics emitted by the variant caller and summarized from the input VCF. This QC class is intended for triage and reproducible filtering. It does not replace manual inspection in Integrative Genomics Viewer (IGV^21–23^ but instead prioritizes variants that warrant review and reduces the likelihood of artifacts. Germline VCF outputs often contain variants supported by minimal evidence, including low alternate read support, mapping artifacts, strand bias, positional bias, and low base quality. A single scalar QUAL value is often insufficient to identify these patterns consistently across samples and cohorts. The QC system, therefore, evaluates multiple orthogonal indicators of evidence quality, including read support, mapping quality, base quality, strand balance, read placement bias, and caller-level support metrics.

The implementation uses FreeBayes style INFO tags together with FORMAT derived depth fields. Bias and artifact related fields include EPP, SAP, ODDS, SRF, SRR, RPL, and RPR^2,11,24^.

To make the rules directly interpretable, the pipeline computes a small set of derived metrics. These measures were intentionally kept simple in order to capture common artifact modes with minimal model dependence and assumptions. Derived metrics computed by the tool:

- QA_ratio = QA / (QA + QR) = A normalized measure of how much base-quality mass supports the alternate allele.
- MQ_diff = MQM – MQMR = A mapping-quality imbalance score; large differences often reflect alignment artifacts preferentially affecting one allele.
- Strand_balance = SRF / (SRF + SRR) = A simple strand symmetry measure for alt-supporting reads.

### QC class definition

The QC framework was designed to be conservative so that variants labeled “Moderate-to-High” are enriched for sites likely to represent true signal. A variant is labeled Low if any of the following failure criteria are met (default thresholds shown):

- Low alternate support: AO < 5

◦ Rationale: Very low alternate counts are disproportionately enriched for errors, especially near indels or in low-complexity regions.
- Low mapping quality for alt reads: MQM < 40

◦ Rationale: Poor mapping quality in the alternate-supporting reads frequently indicates misalignment rather than biology.
- Strong mapping imbalance: MQ_diff ≥ 15

◦ Rationale: When alternate reads map substantially better/worse than reference reads, the locus is often affected by alignment ambiguity.
- Weak caller statistical support: ODDS < 10

◦ Rationale: Low odds/support suggest the caller does not strongly distinguish the variant from the background.
- Very weak base-quality support: QA_ratio < 0.1

◦ Rationale: Indicates that most base-quality evidence supports the reference allele; alternate support may be driven by low-quality bases.
- Excessive bias signals: EPP > 6 or SAP > 6

◦ Rationale: Large bias statistics suggest positional/strand artifacts.
- Read-placement collapse: RPL == 0 or RPR == 0

◦ Rationale: alternate evidence observed only from one side can reflect local misalignment or mapping ambiguity around indels.
- Severe strand imbalance: Strand_balance < 0.20

◦ Rationale: Extreme strand asymmetry is a common signature of technical artifacts.

If any of these criteria are triggered, the variant is marked Low QC. This “fail-fast” logic is deliberate as it prioritizes specificity and interpretability over nuanced scoring. If none of the Low-QC criteria trigger, the variant is labeled Moderate-to-High. This category indicates:

- Read support is adequate.
- Mapping and base-quality evidence are generally consistent.
- Bias metrics are not strongly suggestive of an artifact.

Note: The pipeline groups “Moderate-to-High” together by default because a strict “High” call typically requires manual confirmation (e.g., IGV), which is outside the scope of automated processing^21–23^.

### Adjustability and configuration

QC thresholds are user configurable in order to support different assay types, sequencing depths, and error profiles. Thresholds can be tuned according to study design, including whole genome sequencing, exome sequencing, or targeted panels, as well as specimen type and alignment or calling workflow. In the current implementation, QC logic is encoded through explicit threshold values in the post-processing script. These can be retained as stable defaults and can also be exposed as command-line parameters so that the selected values are captured per run to support traceability and reproducibility. Example parameters include --min_ao, --min_mqm, and - -max_epp, and selected values are captured in run logs to support traceability and reproducibility.

### Practical guidance for interpreting QC_class

“Low QC” should not be interpreted as false by definition. Rather, it indicates that the read evidence shows one or more patterns commonly associated with technical artifacts and should therefore be interpreted cautiously or reviewed manually. Similarly, “Moderate-to-High” does not imply clinical validation. It indicates that the observed read evidence is internally consistent and passes conservative automated checks. In downstream analyses, it is often informative to summarize both all variants and the Moderate-to-High subset, stratify by variant class, and interpret rare variants in the context of both allele frequency and QC status.

### Operational definitions and filtering rules

Filtering used in concordance analyses was defined a priori. SNVs were restricted to sites with single base reference and alternate alleles, and each evaluated site was required to have at least 15 reads in the relevant VCF output. Nonsilent variants were defined from VEP annotations as IMPACT equal to MODERATE or HIGH while excluding synonymous, intronic, upstream, downstream, and UTR consequences. Unique loci were defined from the locus collapsed table using chromosome, position, reference allele, and alternate allele after Step 1 normalization, including decomposition of multiallelic records and left alignment where applicable. Concordance between repeat variant sets was quantified using the Jaccard index on sets of unique loci.

### Demonstration cohort and analysis intent

This dataset included whole-genome sequencing of 5 to 6 single normal human colon crypts and one matched bulk control from each of 21 individuals ranging in age from 10 months to 90 years^17^. The cohort was used to test whether Germline VCF Annotator supports reproducible table generation and rapid QC and concordance review at the cohort scale. Downstream analyses focused on a predefined set of DNA damage response and repair genes listed in Supplemental Table S1. The summaries presented here are intended as usage examples of the tool outputs rather than as a basis for broader conclusions regarding germline DDR architecture in the cohort.

### Statistical analysis

All statistical analyses were performed in RStudio using R version 4.2.2. Data handling and summary table construction used a tidy data workflow that included readr, dplyr, tidyr, stringr, and purrr, whereas standard hypothesis testing used base R stats. Each individual contributed one bulk derived control sample, designated C1, and multiple crypt derived technical repeats, designated T1 through T5 when available. Because crypt samples from the same individual are related observations, individual crypts were not treated as independent samples in patient level comparisons.

For replicate consistency analyses, C1 served as the within patient reference and each crypt derived VCF output was evaluated against C1 using prespecified endpoints derived from the locus collapsed DDR table. These endpoints included total variant rows, higher confidence variant rows defined as QC_class equal to Moderate-to-High, ClinVar annotated rows defined as CLIN_SIG not equal to a dash, and a conservative ClinVar pathogenic or likely pathogenic subset defined as annotations containing pathogenic or likely pathogenic without any accompanying benign or likely benign assertion.

For within patient bulk versus crypt comparisons, the single bulk control sample was compared with the mean of the available crypt samples from the same patient. Reported bulk minus mean crypt differences therefore represent within patient contrasts, and cohort summary values represent the average of those within patient differences across all patients. Paired comparisons were summarized using paired t tests and Wilcoxon signed rank tests.

For age association analyses, patient level summaries were used rather than individual crypt values. Three summaries of the higher confidence DDR variant count, defined as the number of DDR variants classified as Moderate-to-High QC, were evaluated for each patient. These included the bulk control value, the mean crypt value obtained by averaging the available T1 through T5 samples for that patient, and the within patient bulk minus mean crypt difference. Associations with age were quantified using both Pearson and Spearman correlation coefficients.

## RESULTS

### Strategy and key features of the workflow

The motivation for developing the Germline VCF Annotator arose from examining genes in specific biological pathways where germline variation may alter the overall mutation landscape in a whole-genome sequencing study of normal human colon crypts. The Germline VCF Annotator retained explicit reference and alternate alleles and generated two complementary tables, a transcript resolved table and a locus collapsed table, both keyed by genomic position and alleles in human-readable outputs. The workflow also integrated adjustable quality control to summarize depth, mapping quality, strand balance, and related evidence in a consistent tabular format (Fig. 1).

### Transcript resolved and locus collapsed tables were generated at a cohort scale

To establish feasibility for cohort level use, the workflow was benchmarked on 127 germline whole genome sequencing VCF outputs at 30x (Fig. 1). The initial setup phase took approximately 2.6 hours, after which it stabilized at ∼14.2 minutes per VCF. Step 2 completed in 32.8 hours of wall time, producing 11,566,884 transcript-level rows and 1,835,413 locus-collapsed rows (Supplemental Table S2). The workflow produced two complementary outputs at scale, a transcript resolved table containing on the order of ten million annotation rows and a locus collapsed table containing on the order of one to two million unique loci (Fig. 1). These results show that the workflow can process cohort scale genomic datasets and generate both transcript resolved and locus collapsed outputs.

### Locus collapsed output supports rapid concordance assessment across technical repeats

Technical stability was evaluated using within patient repeat VCFs and the locus collapsed output, in which each variant is represented once by a stable locus key defined by Chrom, Pos, Ref, and Alt after normalization. For within-patient comparisons, the same primary bulk sample (C1) was reprocessed across independent crypt-bulk pair analyses, in which the same C1 was paired separately with each available technical repeat crypt sample (for example, C1-T1, C1-T2, and C1-T3). This design allowed the repeated C1 outputs to serve as an internal control for technical stability within each patient, and the representation makes concordance a direct set comparison of loci across samples. Concordance was computed for locus-collapsed DDR SNVs with read depth ≥15x, and that passed the predefined filter set-up described in Methods. Similarity was quantified using the Jaccard index on sets of unique loci defined by Chrom, Pos, Ref, and Alt. Concordance was near perfect across patients, with a median Jaccard index of 1.00 and a minimum of 0.963 (Supplemental Table S3). Discordant loci were enriched for Low QC classifications, indicating that repeat differences were driven predominantly by borderline read evidence rather than instability of germline variant detection. The concordance assessment demonstrated that the workflow produces consistent and reproducible results with repeated independent samples.

### ClinVar annotation facilitates the identification of variants with known clinical significance

The locus collapsed DDR output was next summarized with respect to QC classification and ClinVar^10^ annotations to illustrate the reporting utility of the final tables (summarized in Supplemental Table S2). Across the 127 sample bundles, the DDR locus-collapsed output comprised 1,835,413 variant records. Of these, 149,169 (8.13%) carried a ClinVar clinical significance annotation (CLIN_SIG ≠ “-”). Using a conservative definition, records were retained only when ClinVar contained pathogenic or likely pathogenic assertions, with no accompanying benign or likely benign assertions. Across the 127 sample bundles, 76 sample level locus collapsed records met these criteria (0.004% of all collapsed variants; 0.051% of ClinVar-annotated variants). Because the same locus could be present in more than one technical repeat from the same individual, the 76 records do not represent 76 patient unique loci. Quality stratification indicated that 1,343,664 (73.21%) variants were classified as Moderate-to-High confidence (QC_class = “Moderate-to-High”), indicating that most DDR region calls retained sufficient read evidence for downstream interpretation. After restricting to ClinVar pathogenic or likely pathogenic records of Moderate-to-High QC, 21 sample level records remained for manual review, corresponding to 6 unique loci across the cohort. Together, the ClinVar annotation and QC class fields in the output tables provide a practical basis for prioritizing variants with known clinical significance for targeted review.

### DDR loci showed high concordance across biological repeats in bulk control and mean crypt

To determine whether the applied calling and filtering framework identified any systematic differences between the bulk control sample and the matched crypt samples, locus collapsed DDR variant summaries were compared within each patient (n = 21; Supplemental Tables S4 and S5). For each patient, the single bulk control sample was compared with the mean of the available crypt samples (T1-T5). This was a bulk versus mean crypt comparison within each patient (intra-difference), not a series of pairwise comparisons between the bulk control sample and individual crypt samples (bulk to single crypt difference). Four endpoints were evaluated from the locus collapsed DDR table, including total variants, higher confidence variants, ClinVar annotated variants, and ClinVar pathogenic or likely pathogenic variants.

Intra-differences from all samples were limited, with only small differences (Supplemental Tables S4 and S5). The reported Δ values represent intra-difference within each patient, and mean Δ denotes the average of the intra-differences of all the patients in the cohort. Total variant counts were modestly lower in the bulk control sample (mean Δ = −234.8; −1.62%; paired t-test p = 1.54 × 10⁻⁸; Wilcoxon p = 5.95 × 10⁻^5^), whereas higher confidence variant counts were modestly higher in the bulk control sample (mean Δ = +363.6; +3.47%; paired t-test p = 1.58 × 10⁻¹⁶; Wilcoxon p = 9.54 × 10⁻⁷). ClinVar annotated counts showed a similarly small decrease in the bulk control sample (mean Δ = −12.1; −1.05%; paired t-test p = 1.98 × 10⁻⁵; Wilcoxon p = 2.29 × 10⁻4), whereas ClinVar pathogenic or likely pathogenic counts did not differ between the bulk control sample and the mean crypt summary (mean Δ = +0.08; paired t-test p = 0.202; Wilcoxon p = 0.248). Taken together, these results indicate that bulk control and mean crypt DDR burden were highly concordant across biological repeats, with only modest differences across the endpoints examined.

### Higher confidence DDR loci showed no age associated trend

Age associations were evaluated across the 21 patients (inter-difference) using three patient level values derived from Supplemental Table S4 and summarized in Supplemental Table S5, the bulk control summary (C1_MH), the mean crypt summary obtained by averaging the available T1 through T5 samples within each patient (Mean_T1 to T5_MH), and the within patient bulk minus mean crypt difference (intra-difference; Delta_MH_C1_minus_Tmean). Age was not associated with higher confidence DDR burden in the bulk control sample (Pearson r = −0.263, p = 0.249; Spearman ρ = −0.283, p = 0.214). Likewise, no age association was found with higher confidence DDR burden in the mean crypt summary (Pearson r = −0.224, p = 0.328; Spearman ρ = −0.310, p = 0.171), or in the intra-difference summary (Pearson r = −0.281, p = 0.217; Spearman ρ = −0.339, p = 0.133; Table S5). These findings showed very little inter-difference among the 21 patients, indicating no age association of higher confidence DDR burden in the normal human colon crypts.

### IGV assessment of QC prioritized ClinVar flagged DDR loci

The 21 ClinVar flagged DDR records classified as Moderate-to-High QC were reviewed at the read evidence level using IGV to contextualize discordance across within patient technical repeats and to evaluate technical validity (Supplemental Table S6; Fig. 2). Loci with stable support across repeats were generally compatible with true underlying germline variation, whereas discordant observations were concentrated among sites with sparse alternate support, strand asymmetry, or locally complex sequence context.

**Fig. 2.**
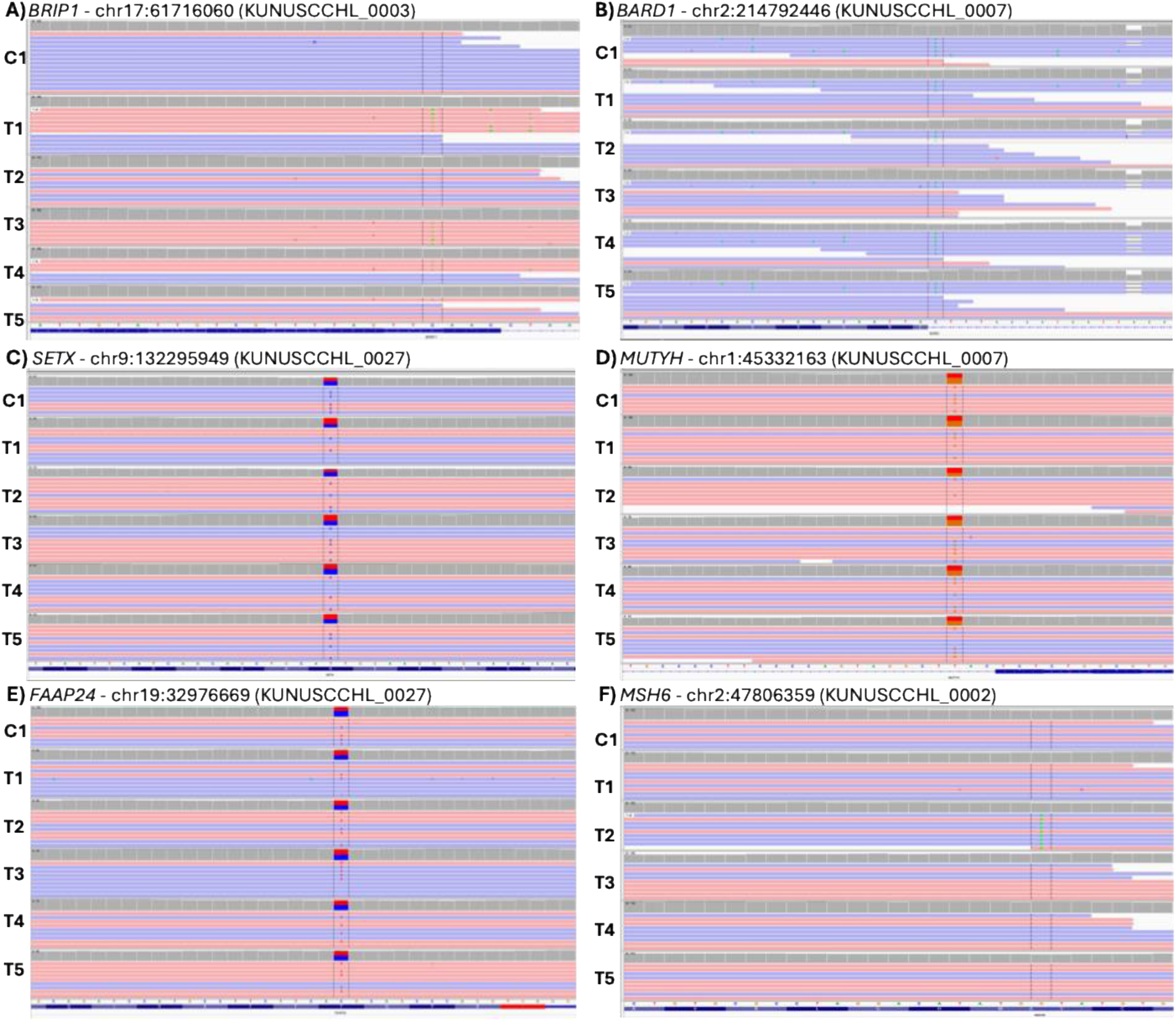
IGV read-level evidence for ClinVar-flagged DDR loci. Integrative Genomics Viewer (IGV) snapshots are shown for candidate DDR loci highlighted in Table S6 that were manually reviewed for variant calling accuracy. Each panel displays aligned reads and coverage across the corresponding sample tracks (bulk-derived C1 and, where available, technical repeats T1–T5) at the candidate locus in hg38 coordinates; the vertical guide marks the site of interest. Panels illustrate representative read-evidence patterns that informed conservative interpretation of ClinVar-flagged loci: (A) BRCA1 interacting helicase 1 (*BRIP1*), (B) BRCA1-associated RING domain 1 (*BARD1*), (C) senataxin (*SETX*), (D) MUTYH DNA glycosylase (*MUTYH*), (E) Fanconi anemia-associated protein 24 (*FAAP24*), and (F) MutS homolog 6 (*MSH6*). The loci information is summarized in Table S6.

This pattern was evident in *BRIP1* (Fig. 2A) and *BARD1* (Fig. 2B), where the primary limitation was the underlying technical evidence. In both genes, the alternate base was observed only in reads from one direction, with no supporting reads from the opposite direction, raising the possibility that the signal was influenced by strand bias. In addition, the presence of an indel near *BARD1* may suggest a possibility of a local alignment complexity rather than a single-nucleotide event. By contrast, IGV supported genuine underlying variation at several loci, including *SETX* (Fig. 2C), *MUTYH* (Fig. 2D), and *FAAP24* (Fig. 2E), although ClinVar interpretations across these loci were variable, ranging from conflicting pathogenicity classifications to likely pathogenic assertions.

*MSH6* showed a distinct pattern (Fig. 2F). The locus was annotated in ClinVar as likely pathogenic. In patient 0002 crypt T2, the variant was supported by 9 alternate A reads and 29 reference G reads. However, it was not detected in the matched bulk sample or in the remaining crypt repeats from the same individual. This distribution was inconsistent with a germline variant in the patient and supported the possibility of a crypt restricted variant.

Taken together, QC prioritized IGV review distinguished technically credible loci from observations limited by sparse support, strand asymmetry, or local alignment complexity. These findings show how the workflow can organize flagged loci for manageable targeted IGV review and support interpretation of technically credible germline and candidate somatic variants.

## DISCUSSION

The colon crypt cohort provided a biologically relevant demonstration of how the Germline VCF Annotator can support variant interpretation by generating an organized list of variants in a group of genes from a chosen pathway for human-manageable final review to determine their biological relevance. The workflow processed millions of variants and generated stable human readable tables, supported conservative QC based prioritization, and facilitated targeted IGV review of ClinVar flagged variants within a gene set of interest. Among the final six loci, *SETX, FAAP24*, and *MUTYH* illustrated variants with technically credible evidence that warranted further study-specific evaluation. Two other loci, *BRIP1* and *BARD1,* illustrated observations whose support was weakened by sparse read evidence, strand asymmetry, or local alignment complexity. *MSH6* showed a distinct pattern, appearing in a single crypt but not in the matched bulk or other crypt repeats from the same patient. In addition to identifying potentially biologically meaningful germline variants, the workflow originally intended to achieve, the workflow can also surface candidate somatic events when appropriate controls are available. The biological relevance of any such locus identified by the workflow can then be determined by the molecular question under study and the surrounding evidence. It is important to note that the annotator itself does not establish biological significance; it only provides an effective framework for identifying, organizing, and prioritizing candidate variants for human-manageable review and downstream interpretation. The exercise also showed that ClinVar based filtering alone is insufficient for final interpretation without secondary review of the underlying read evidence. The workflow presented here provides a practical way to reduce the candidate set to a manageable number of higher-confidence loci for targeted IGV review, thereby improving confidence in retained variants and reducing false-positive carry-through.

This study also demonstrated that the workflow benefits from a modular structure that supports reproducibility and scale. Separating VCF normalization and annotation from downstream extraction and QC allows the annotated TSV layer to be reused for multiple purposes, including gene focused extraction, cohort level summaries, and concordance benchmarking, without rerunning the more computationally expensive upstream steps. This organization also extends the utility of the framework beyond the present application. Although the motivating use case centered on curated DDR genes in normal colon crypts, the same design can be applied to other predefined gene groups or to more general germline reporting tasks in research and translational settings. The implementation presented here used FreeBayes-derived germline VCF outputs, but the downstream workflow can also be applied to germline VCFs generated by other callers, provided that the input VCF files retain the standard fields in the standard VCF output of all variant callers.

Several limitations should be acknowledged. First, the QC classification is intentionally rules based and heuristic. It summarizes interpretable read evidence patterns for transparent triage, but borderline loci may shift between classes according to sequencing depth, alignment context, or caller specific behavior. Second, annotation completeness and field naming depend on the reference build, transcript set, and annotation resource versions. Third, ClinVar labels are not equivalent to genotype level support within a specific dataset under fixed thresholds, and clinically flagged loci still require evaluation in the context of read evidence and repeat structure. Fourth, the present workflow is focused on small germline variants and does not address structural variation, copy number change, or low level mosaicism, each of which requires distinct calling strategies and evidence models.

Future development should focus on increasing interoperability and reporting flexibility. One useful extension would be an optional export to Mutation Annotation Format (MAF) to support compatibility with established tabular mutation pipelines and downstream analytic frameworks. Additional transcript prioritization layers, such as MANE Select or canonical transcript views, could also be incorporated while preserving the full transcript resolved output. Further QC summary modules, including per sample QC distributions, stratification by variant class and allele frequency, and automated concordance summaries across repeats, would strengthen cohort level benchmarking and support broader comparative analyses.

In summary, Germline VCF Annotator provides a practical and transparent approach to convert germline VCF outputs into stable, human readable tables that preserve both transcript aware biological annotation and evidence context. By directly addressing transcript multiplicity, explicit allele identity, and evidence based QC grading, the workflow supports reproducible germline reporting, conservative variant interpretation, and biologically focused exploratory analysis in settings where locus level clarity is essential.

## CODE AVAILABILITY

Germline VCF Annotator is publicly available (source code) on GitHub (https://github.com/Z-Mano/Germline-VCF-Annotator), including the complete two-step workflow scripts, documentation, and example configuration files. The software is licensed for academic and research use only (© 2026 Zarko Manojlovic and Chih-Lin Hsieh; see LICENSE). Releases are versioned and tagged in the repository; to support reproducibility, users should cite the specific release tag and, when relevant, the corresponding commit hash. A changelog and release notes provide a stable record of updates over time.

## ACKNOWLEDGMENTS

The author thanks Drs. Chih-Lin Hsieh, Michael Lieber, and Yong-Hwee Eddie Loh for their valuable input, collaboration, and access to the data^17^ used in this study. This work was supported by the National Institute on Aging (R01 AG067615) and the Catherine and Joseph Aresty Endowment.

## SUPPLEMENTARY TABLES

**Supplemental Table S1. DNA Damage Response and Repair (DDR) gene set used for targeted analyses.** Curated list of the 160 DDR genes evaluated in this study, organized by functional pathway.

**Supplemental Table S2. Per-sample summary of DDR-region variant burden, quality stratification, and ClinVar annotation.** Per-sample summary of locus-collapsed variant counts within the predefined DDR gene set, including the bulk control sample (C1) and crypt repeat samples (T1 through T5), together with subject metadata. This table provides a compact overview of variant burden, higher confidence calls, and ClinVar annotated variants across the analyzed samples. Column definitions and abbreviations are provided in the table footnote.

**Supplemental Table S3. Bulk (C1) technical replicate concordance of nonsilent DDR loci (FreeBayes; locus-collapsed).** Summary of within-patient concordance across the bulk control sample and available crypt repeats for nonsilent DDR loci after locus collapsing and filtering as described in Methods. Reported fields include repeat- specific locus counts, union and intersection counts, pairwise Jaccard similarity values, mean and minimum Jaccard statistics, and the corresponding totals for nonsilent entries classified as Moderate-to-High or Low QC. For patient KUNUSCCLH_0012, T2 failed sequencing and was excluded. A replacement sample, T6, was generated, but the primary summaries in this table were restricted to the prespecified repeat set T1 through T3 for consistency across patients. As a result, n_repeats = 2 for this individual.

**Table S4. Patient-level bulk control and mean crypt DDR variant summaries used for within-patient comparisons (intra-difference) and age association analyses (inter-difference).** Intra-differences between the baseline bulk callset (C1) and the mean of available technical repeats (n_T_repeats indicates the number of crypts present) for each of the 21 patients are summarized. For each of the 21 patients, locus-collapsed variant summaries across the predefined DDR gene set are reported for the bulk control sample (C1) and for the mean of the available crypt repeat samples within the prespecified repeat set T1 through T5. Reported fields include total variant counts, higher confidence variant counts, ClinVar annotated variant counts, and ClinVar pathogenic or likely pathogenic variant counts, together with the corresponding bulk minus mean crypt differences and percentage differences. Mean_T1 to T5 denotes the mean calculated across the available crypt repeat samples for that patient and does not represent a comparison between C1 and T5 alone. n_T_repeats indicates the number of crypt repeat samples contributing to the mean summary for that patient. Age is included to support the age association analyses summarized in Supplemental Table S5. For patient KUNUSCCLH_0012, crypt T2 failed sequencing and was excluded. A replacement sample, T6, was generated, but the primary summaries in this table were restricted to the prespecified repeat set T1 through T5 for consistency across patients. As a result, n_T_repeats = 4 for this individual.

**Supplemental Table S5. Statistical summary of within-patient bulk versus mean crypt DDR comparisons (intra-difference) and age association analyses (inter-difference).** For each of the 21 patients, the single bulk control sample (C1) was compared with the mean of the available crypt samples within the prespecified repeat set T1 through T5. In the first section, summary statistics are reported for four endpoints derived from the locus-collapsed DDR table, including total variants, higher-confidence variants, ClinVar annotated variants, and ClinVar pathogenic or likely pathogenic variants. Mean Δ denotes the cohort mean of the intra-difference calculated as bulk minus mean crypt, and percent difference is expressed relative to the mean crypt value. Paired t test and Wilcoxon signed rank p values summarize the within-patient comparison across the 21 patients. In the second section, age association statistics are reported for three patient-level higher-confidence DDR summaries, the bulk control value (C1_MH), the mean crypt value computed from the available T1 through T5 samples for that patient (Mean_T1 to T5_MH), and the within-patient bulk minus mean crypt difference (Delta_MH_C1_minus_Tmean). Pearson and Spearman correlation statistics were calculated across the 21 patients using AGE from Supplemental Table S4 paired with these patient-level higher-confidence DDR summaries.

**Supplemental Table S6. High-confidence ClinVar-flagged DDR variants identified from the human normal colon crypt study.** List of DDR variants retained for targeted review after filtering for ClinVar annotation and Moderate to High QC support. For each retained variant instance, the table reports sample identifiers, genomic coordinates, gene symbol, transcript level annotation, ClinVar interpretation, population frequency, and selected FreeBayes read evidence fields used to support QC classification. Recurrent entries across samples from the same individual reflect repeat level recovery of the same locus, whereas single sample occurrences indicate loci detected in only a subset of repeats.

## REFERENCES

1. Danecek P, Auton A, Abecasis G, et al. The variant call format and VCFtools. Bioinformatics. 2011;27(15):2156–2158.

2. Garrison E, Marth, Gabor. Haplotype-based variant detection from short-read sequencing. arXiv. 2012;arXiv:1207.3907.

3. Richards S, Aziz N, Bale S, et al. Standards and guidelines for the interpretation of sequence variants: a joint consensus recommendation of the American College of Medical Genetics and Genomics and the Association for Molecular Pathology. Genet Med. 2015;17(5):405–424.

4. Cingolani P, Platts A, Wang le L, et al. A program for annotating and predicting the effects of single nucleotide polymorphisms, SnpEff: SNPs in the genome of Drosophila melanogaster strain w1118; iso-2; iso-3. Fly (Austin). 2012;6(2):80–92.

5. McLaren W, Gil L, Hunt SE, et al. The Ensembl Variant Effect Predictor. Genome Biol. 2016;17(1):122.

6. McCarthy DJ, Humburg P, Kanapin A, et al. Choice of transcripts and software has a large effect on variant annotation. Genome Med. 2014;6(3):26.

7. Morales J, Pujar S, Loveland JE, et al. A joint NCBI and EMBL-EBI transcript set for clinical genomics and research. Nature. 2022;604(7905):310–315.

8. Yen JL, Garcia S, Montana A, et al. A variant by any name: quantifying annotation discordance across tools and clinical databases. Genome Med. 2017;9(1):7.

9. Karczewski KJ, Francioli LC, Tiao G, et al. Author Correction: The mutational constraint spectrum quantified from variation in 141,456 humans. Nature. 2021;590(7846):E53.

10. Landrum MJ, Lee JM, Benson M, et al. ClinVar: improving access to variant interpretations and supporting evidence. Nucleic Acids Res. 2018;46(D1):D1062–D1067.

11. McKenna A, Hanna M, Banks E, et al. The Genome Analysis Toolkit: a MapReduce framework for analyzing next-generation DNA sequencing data. Genome Res. 2010;20(9):1297–1303.

12. Kandoth C. mskcc/vcf2maf: vcf2maf v1.6. Zenodo. 2020.

13. Cagan A, Baez-Ortega A, Brzozowska N, et al. Somatic mutation rates scale with lifespan across mammals. Nature. 2022;604(7906):517–524.

14. Lee-Six H, Olafsson S, Ellis P, et al. The landscape of somatic mutation in normal colorectal epithelial cells. Nature. 2019;574(7779):532–537.

15. Hsieh CL, Manojlovic Z, Okitsu T, et al. Analysis of Naturally Occurring Somatic Insertions in the Human Genome. bioRxiv. 2025.

16. Manojlovic Z, Okitsu C, Okitsu T, et al. High-depth Whole Genome Sequencing of Single Human Colon Crypts Uncovers New View on Crypt Clonality. bioRxiv. 2025.

17. Manojlovic Z, Wlodarczyk J, Okitsu C, et al. Construction of high coverage whole-genome sequencing libraries from single colon crypts without DNA extraction or whole-genome amplification. BMC Res Notes. 2023;16(1):66.

18. Chatterjee N, Walker GC. Mechanisms of DNA damage, repair, and mutagenesis. Environ Mol Mutagen. 2017;58(5):235–263.

19. Schumacher B, Pothof J, Vijg J, Hoeijmakers JHJ. The central role of DNA damage in the ageing process. Nature. 2021;592(7856):695–703.

20. Danecek P, Bonfield JK, Liddle J, et al. Twelve years of SAMtools and BCFtools. Gigascience. 2021;10(2).

21. Robinson JT, Thorvaldsdottir H, Wenger AM, Zehir A, Mesirov JP. Variant Review with the Integrative Genomics Viewer. Cancer Res. 2017;77(21):e31–e34.

22. Robinson JT, Thorvaldsdottir H, Winckler W, et al. Integrative genomics viewer. Nat Biotechnol. 2011;29(1):24–26.

23. Thorvaldsdottir H, Robinson JT, Mesirov JP. Integrative Genomics Viewer (IGV): high-performance genomics data visualization and exploration. Brief Bioinform. 2013;14(2):178–192.

24. DePristo MA, Banks E, Poplin R, et al. A framework for variation discovery and genotyping using next-generation DNA sequencing data. Nat Genet. 2011;43(5):491–498.

